# Evolution of individual variation in a competitive trait: a theoretical analysis

**DOI:** 10.1101/2023.12.06.570103

**Authors:** Klaus Reinhold, Lukas Eigentler, David W. Kikuchi

**Affiliations:** Department of Evolutionary Biology, Bielefeld University, Germany; Department of Integrative Biology, Oregon State University, OR, USA

**Keywords:** maintenance of variation, standing variation, negative frequency dependence, individual based model, resource distribution, temporal fluctuations

## Abstract

When competitive traits are costly, negative frequency-dependence can maintain genetic variance. Most theoretical studies examining this question assume binary polymorphisms, yet most trait variation in wild populations is continuous. We propose that continuous trait variation can result from continuous variation in resource quality and that the shape of the resource distribution determines trait maintenance. We used an individual-based model to test which conditions favour the stable maintenance of variation and which cause temporal fluctuations in trait values. This approach, inspired by contrasting outcomes of previous studies, clearly showed the decisive role played by the shape of resource distributions. Under extreme conditions, like the absence of resource variation or with very scarce resources for weak competitors, traits evolved to a single non-competitive or highly competitive strategy, respectively. Some distributions led to strong temporal fluctuations on trait values, whereas others led to the maintenance of large standing variation in competitive traits together with stable mean trait values. Our results thus explain the contradicting outcomes of previous theoretical studies and at the same time provide hypotheses to explain the maintenance of genetic variation and individual differences. We also suggest how the proposed effects of resource variation on trait maintenance can be tested empirically.

## Introduction

High values for evolvability and heritability have frequently been found for traits that are supposedly under strong selection (e.g., Houle 1992, Merilä & Sheldon 2000). This is surprising, as selection should tend to erode standing genetic variation according to Fisher’s fundamental theorem (Grafen 2015). There are several theoretical explanations for such findings. Temporal or spatial fluctuations in selection could maintain genetic variation in populations, as traits that are under positive selection in some environments can be selected against in other environments (Reinhold 2001, Svardal et al. 2015, Radwan et al. 2016). Another widely discussed possibility is the presence of negative frequency-dependent selection, meaning that the direction of selection changes from positive to negative with increasing frequencies of phenotypes (Trotter & Spencer 2007). Such negative frequency-dependent selection has, for example, been invoked in the evolution of variation regarding the resistance towards parasites or diseases. Bacteria and parasites tend to be better adapted to more frequent genotypes and thus selection can favour rarer host genotypes (Hamilton & Zuk 1982). Another source of negative frequency-dependent selection may be competition for resources. Theoretical examinations of the evolution of costly investments into competition have shown that negative frequency-dependent selection can lead to the maintenance of genetic variation under many environmental conditions (Wolf et al. 2007, 2008; Baldauf et al. 2014, Kikuchi & Reinhold 2021). According to such theoretical results, negative frequency-dependent selection is especially likely if individuals compete most strongly with individuals in the population that have similar traits, and less with dissimilar individuals.

The vast majority of theoretical studies on frequency-dependent selection and its role in maintaining genetic variation assume a small discrete number of genotypes. Often, only two or three alleles are assumed to produce a limited number of phenotypes (Sinervo & Lively 1996, McElreath & Strimling 2006, Wolf et al. 2007, 2008, Harris et al. 2008, Broom et al. 2018, Christie et al. 2018). These simple models of negative frequency-dependent selection are nevertheless very helpful in giving criteria for the maintenance or erosion of genetic variation, for example in small populations, under inbreeding, with temporal or spatial fluctuations in selection, or when overdominance in fitness is present. However, with these simplified models, it is very hard to give quantitative predictions of the extent of genetic or phenotypic variation that can be maintained and how this depends on environmental conditions. A few studies have allowed a continuum of phenotypes to occur simultaneously in populations. In one of these, Baldauf et al. (2014) used individual-based simulations to show that dynamic fluctuations in the value of traits frequently occur when individuals with costly competitive traits compete for resources that vary in quality. Here, “costly” means that resources that could otherwise be utilised for offspring production are invested into competitive ability. In most of their analyses, Baldauf et al. (2014) assumed that there were only two types of resources: good resources that were rare, and poor resources that were unlimited. Under these conditions, recurring fluctuations in mean trait values resulted. These fluctuations were caused by an initial arms race in the development of competitive traits resulting in extreme trait values, which subsequently allowed the invasion of less competitive trait values. In contrast to Baldauf et al. (2014), another study showed that under broad and realistic conditions of costly competition, temporally stable polymorphisms of many genotypes were often maintained (Kikuchi & Reinhold 2021). Even though there are many similarities between these two approaches, the differences in the stability of the outcomes are striking. Baldauf et al. (2014) obtained dynamic fluctuations in mean trait value assuming binary or uniform distributions of resource quality. In contrast, Kikuchi & Reinhold (2021) observed stable polymorphisms when they assumed resource quality to be distributed like the right half of a normal distribution. However, Kikuchi & Reinhold (2021) also observed temporal fluctuations under uniform resource distributions. This leads to the hypothesis that resource distributions have a large influence on the evolution of costly competitive adaptations.

In the current study, we examine the influence of the shape of resource distributions on the evolution of costly investments into competition and on temporal trait stability. Similar to Baldauf et al. (2014), we examine the evolution of costly competitive traits in an asexually reproducing population, but use several different resource distributions to examine under which conditions temporally stable or fluctuating polymorphisms are produced. Our main aim is to test whether the fluctuating trait values observed by Baldauf et al. (2014) and the stable maintenance of large trait variance observed by Kikuchi & Reinhold (2021) rely on the specific conditions assumed in their models or are likely to occur under a wide array of resource distributions, i.e., ecological conditions.

## Methods

We used an individual-based model to test the evolutionary dynamics of costly competitive traits that determine the quality of resources an individual is able to obtain. We denote the trait by *c* ≥ 0, where *c* = 0 corresponds to no investment into competitive traits, and higher *c* values signify higher energetic investments. To facilitate direct comparison, we assumed the same trade-off between competitive traits and reproduction as Baldauf et al. (2014). We assumed that any individual that attains a resource of quality *r*, can only invest a proportion of 1 − *c* into reproduction because remaining resources are required to develop and maintain their competitive ability.

Note that we impose no upper bound on the trait value *c* and, in principle, allow *c* ≥ 1 to occur through mutation (see below). In such cases, we assume that no energy can be allocated for reproduction which immediately leads to selection of *c* < 1. Moreover, we show below that *c* = 1 − *d*, where *d* denotes the ratio of the resource quality that can be obtained without investment into competitiveness (*c* = 0) to the quality of the best available resources, forms a natural upper bound for the competitive trait.

We used a fixed population size of 10000 individuals. In the first generation, we always started our simulations by randomly choosing a value of *c* on the interval [0,1] for each individual. Independent of the resource distribution used, for each individual and each generation, we varied the value of *c* by adding random values from a normal distribution with SD of 0.01 to find the ‘expressed’ value ĉ, so that there were slight deviations from the ‘genetic’ value of *c*. This is similar to the approach of Baldauf et al. (2014) and smoothens the curve of the relationship between genotype and fitness. It can be interpreted as the effect of developmental noise, mimicking chance effects on the phenotype when individuals with very similar genetic traits compete. We assumed that the individual that expressed the most competitive trait obtained the best available resource and so on, such that the individual with the lowest value of ĉ obtained the least valuable resource. For every generation, 10000 resources were drawn randomly from the chosen distribution. We chose the product of *r*, the resource quality obtained, and the proportion of resources not invested into the competitive trait (1 − *c*), to calculate the reproductive fitness of each individual strategy. We assumed asexual reproduction. Using these fitness values, we chose 10000 offspring using a lottery so that individuals were expected to reproduce in proportion to their fitness. For the offspring, we assumed like Baldauf et al. (2014) a mutation rate of 0.01 for the genetic value of *c* (i.e., 1% of the offspring mutated), and a normally distributed effect of the mutation with SD 0.1. If mutations led to *c* values smaller than 0, we set them to 0 (i.e., there was a lower bound for *c* that represented no development of any competitive traits). We recorded the population mean value of *c*, the SD of the distribution of trait values at generation 1000, and the minimum and maximum mean value of *c* between generations 1000 and 2000. For each specific condition we ran 100 independent simulations, which was sufficient to yield consistent patterns.

To represent various types of resource distributions, we used four simplistic linear relationships (A to D) and two non-linear ones (E and F), and without loss of generality assumed that the best resources have quality 1 and the worst resources have quality *d*, with *d* varying from 0 to 1 (see Fig. 1). We ordered the distributions according to an increase in the availability of low-quality resources from A to F. The distribution with the lowest frequency of poor-quality resources assumed a linear increase in the availability of resources between *d* and 1 (distribution A). To allow a comparison with mid-centred unimodal distributions, we used distribution B, where resources with value (1 + *d*)/2 have twice the frequency compared to a uniform distribution and resource values towards *d* and 1 linearly approach frequencies of 0 (see figure 1). We also used the uniform distribution (distribution C). Distribution D is a linear distribution with more poor resources than good ones, where resource abundance decreases linearly from resources with quality *d* that have an abundance of twice the level of the uniform distribution C, to resources of quality 1 that have an abundance of 0 (and thus constitutes a mirror image of distribution A). As we (Kikuchi & Reinhold 2021) did not observe trait fluctuations in a model similar to the one by Baldauf et al. (2014) when using the right half of a normal distribution, we also used such a distribution (distribution E), defining it to be the right half of a normal distribution with mean *d* and SD of (1 − *d*)/3, and truncating it at +3SD to avoid resource values above unity (Fig. 1). Finally, we used a power law (distribution F) because we can analytically show (see appendix) that it leads to a uniform distribution of the competitive trait value.

**Figure 1.**
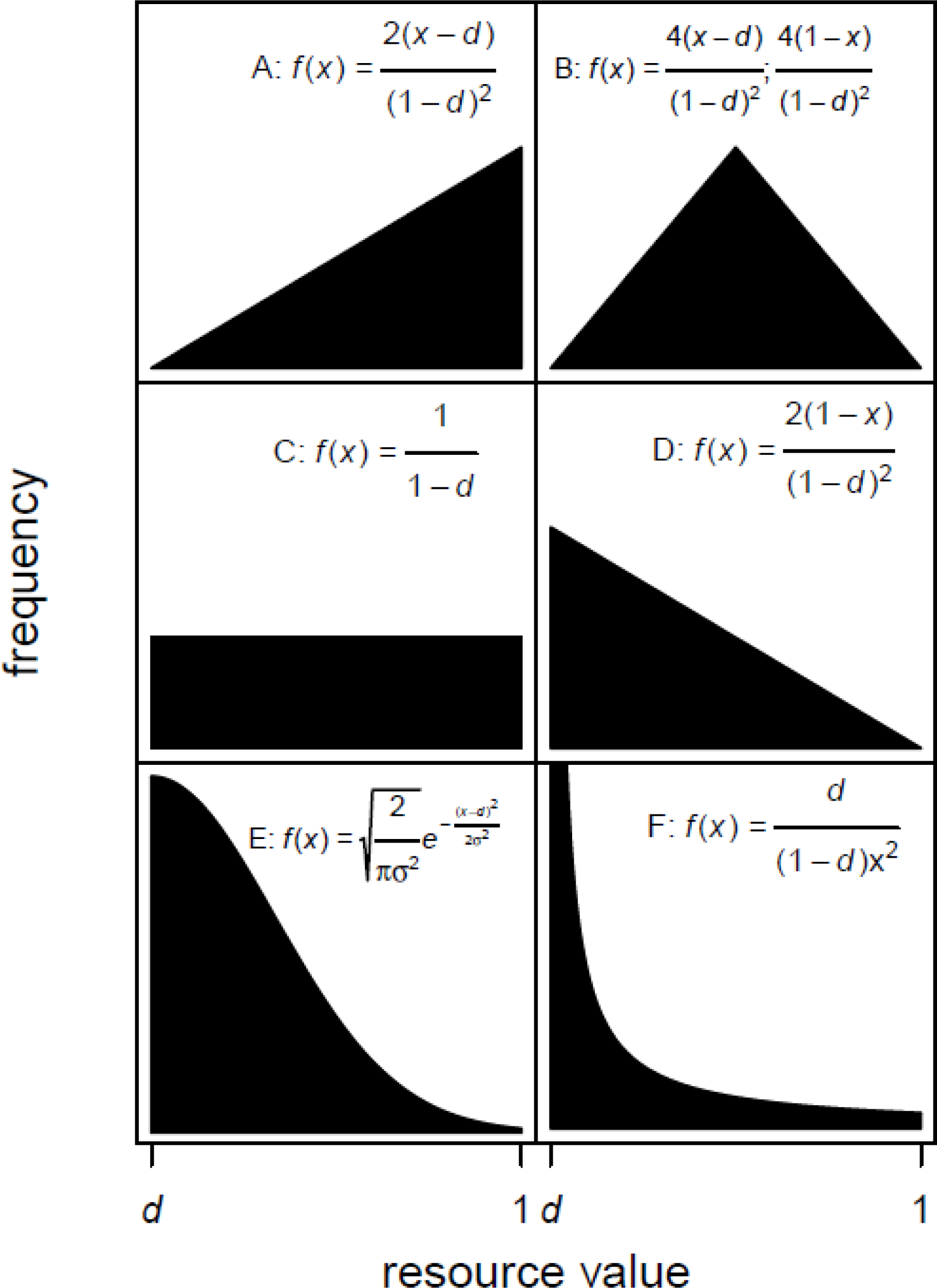
Depiction and description of the resource distributions used in our simulations. The resource values are given on the x-axis and the relative probability densities are given on the y-axis. Equations for the probability density are superimposed within each panel. A. Linear increasing. B. Linear mid-centered. The first equation applies for 0 ≤ x ≤ 0.5, and the second applies for 0.5 < x ≤ 1. C. Uniform distribution. D. Linear increasing. E. Truncated right half of normal distribution. F. Power law distribution.

## Results

The resource distributions, as well as the ratio between the best and the worst resources (i.e., the value of *d*), had a strong influence on the outcome of our simulations. This dependence occurred for all response variables: the average *c*-value (proportion of resources invested into competitive ability) at generation 1000, the range and fluctuation of mean *c*-values between generations 1000 and 2000, and the standing variation of *c* in generation 1000. We report our results for these three metrics separately in the following paragraphs.

### Evolution of average investment into competition

The mean trait value *c* observed at generation 1000 is strongly influenced by the value of *d* and the resource distribution (Fig. 2). Simulations with *d* = 0.01 resulted in an evolutionary stable value of *c* between 0.94 and 0.99 for all used resource distributions except the power law distribution (F). For resource distribution F, we observed markedly lower values of *c*, around 0.5 (see appendix for an explanation why this fits the expectation). As expected, the simulations with *d* = 1, i.e., in the absence of resource variation, all resulted in *c* evolving towards 0 and its value only remained slightly above this due to a mutation-selection balance. At intermediate values of *d*, there are noteworthy differences among the distributions in the mean value of *c*. For intermediate values of *d*, and the linear resource distributions with few low-quality resources (distributions A to C), the observed mean values of *c*, at generation 1000, decreased approximately proportional to 1 − *d* (see Fig. 2). For two of the resource distributions with a majority of low-quality resources (distributions D and E), mean *c* initially decreased more steeply with increasing *d*. For the resource distribution based on a power-law (distribution F), mean *c* decreased linearly and was always very close to (1 − *d*)/2.

**Figure 2.**
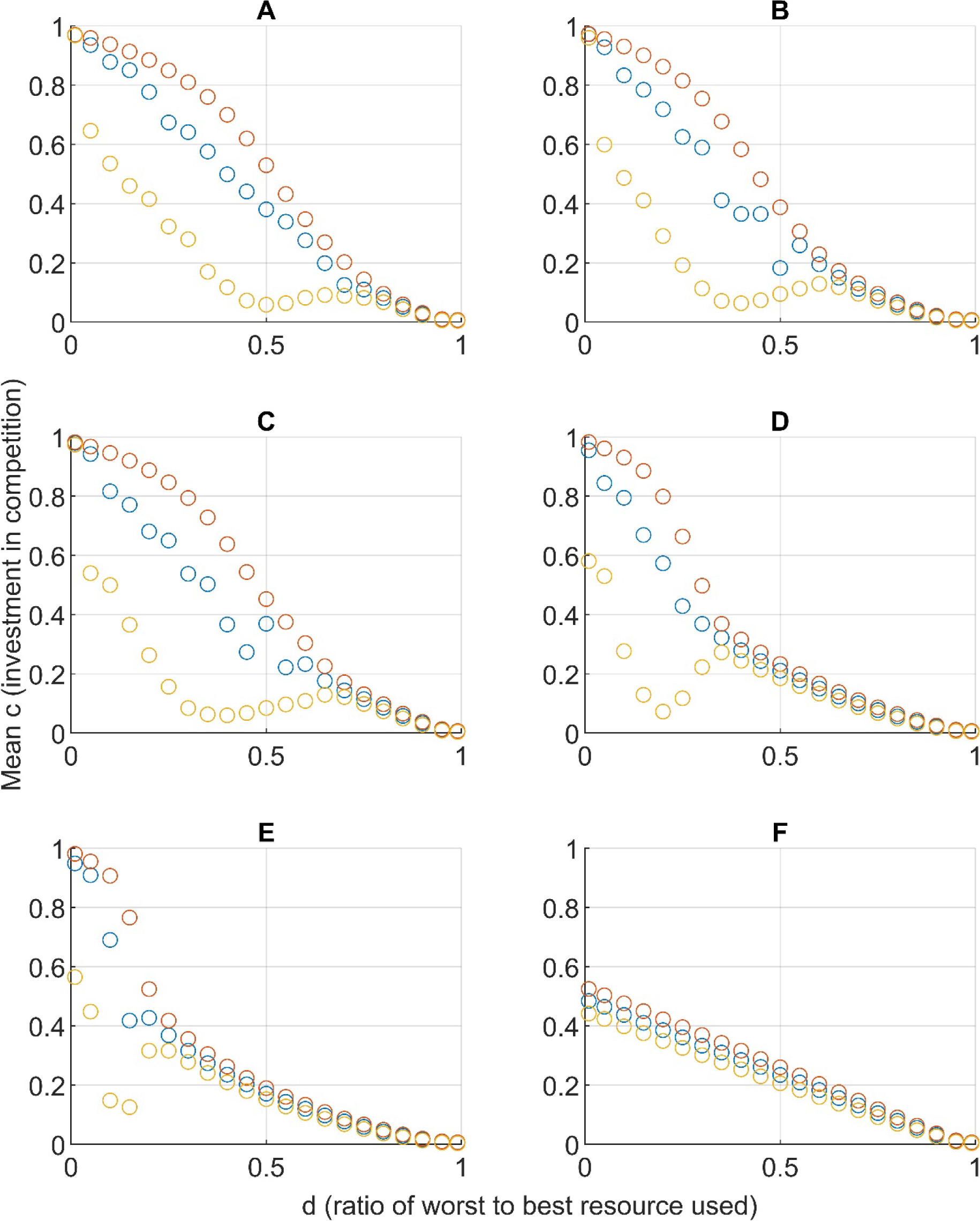
Summary of simulation results dependent on the assumed resource distributions A to F and *d*, the assumed ratio between the worst and the best resource. Depicted are the mean values of the competitive trait *c*, at generation 1000 (blue dots), as well as the range of temporal fluctuations given as the average minimum (yellow dots) and the average maximum (orange dots) between generations 1000 and 2000. Large vertical differences between red and yellow dots indicate strong temporal variations. All averages are calculated from 100 independent repetitions.

### Occurrence of strong temporal fluctuations

For intermediate values of *d*, all resource distributions except the power-law distribution (distribution F) regularly resulted in strong temporal fluctuations of the mean competitive trait between generations 1000 and 2000, as can be seen by the large range between the average minimum and the average maximum for this interval (Fig. 2). Such strong and quick temporal fluctuations occurred for these distributions with values of *d* clearly larger than 0 and up to a certain value of *d*. Above this threshold, the strong temporal fluctuations disappeared (see Fig. 3 for an example on both sides of this threshold value). The threshold was negatively correlated with the proportion of low-quality resources, and thus decreased from distribution A to distribution E. However, such strong temporal fluctuations were absent for resource distribution F, in agreement with our analytical prediction (appendix).

**Figure 3.**
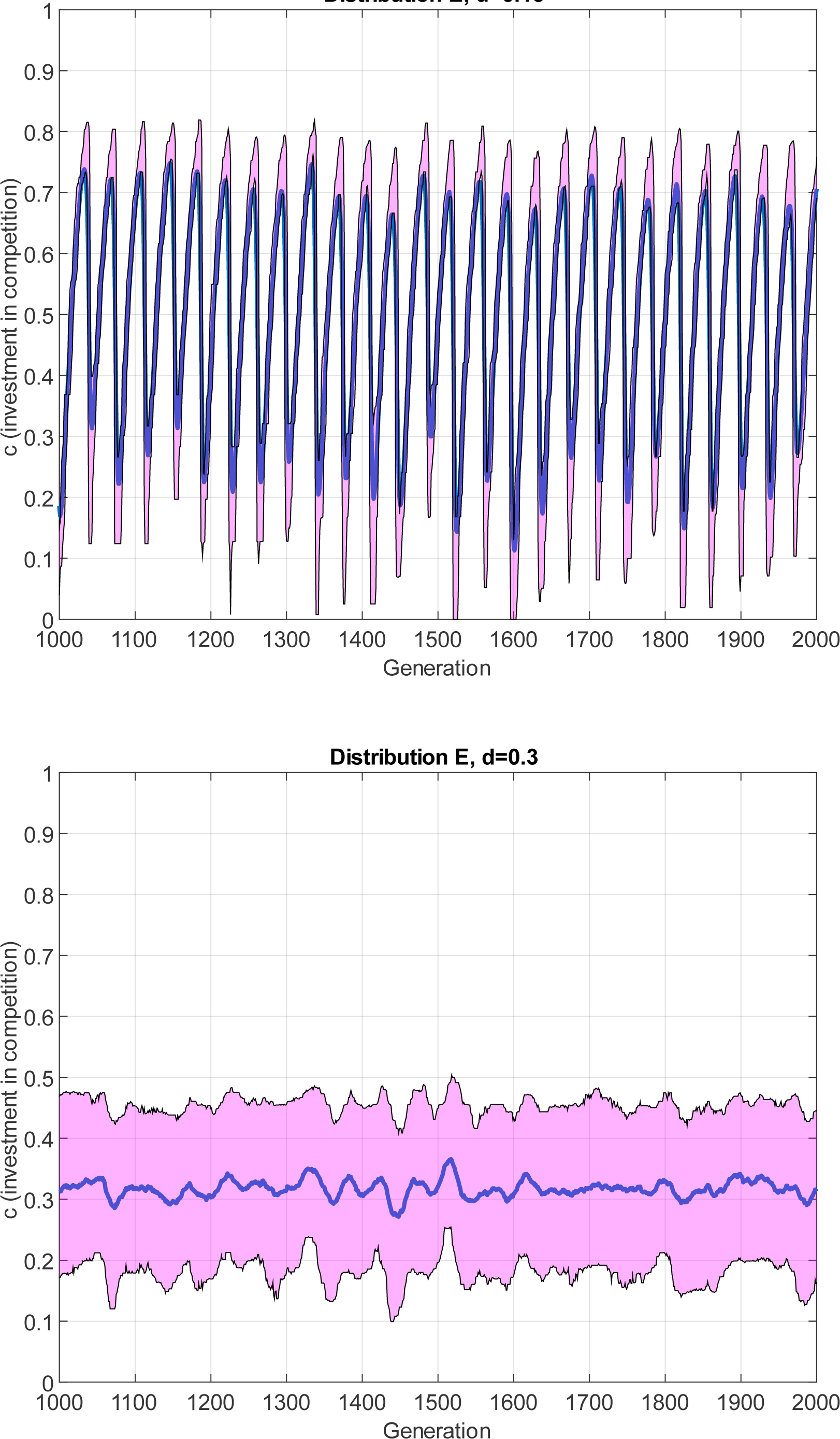
Exemplary results of simulations for the distribution E. For a, the value of d was set to 0.15 and in b, it was set at 0.4. Depicted are the temporal variation of the average c values (blue) as well as the interquartile range (rose) as an estimate for the extent of standing genetic variation.

### Maintenance of standing genetic variation of competitive trait

Some maintenance of standing genetic variation in competitive trait values was evident in all our simulations (see figure 4). Very low standing variation, likely caused by mutation selection balance, only occurred when *d* was assumed to be close to 1 or 0. The amount of standing variation was high for all distributions whenever strong temporal fluctuations were absent. Under these conditions, the standing variation approached the expected maximal value, given the assumption that all competitive trait values *c* ≤ (1 − *d*) are present in equal frequencies (with such a uniform distribution of trait values, the SD of the variation in genetic traits is *SD* = (1 − *d*)/√12). Higher trait values are not expected to persist, as the fitness of the trait with low competitiveness would be at least *d*, and thus higher than the fitness of any trait investing stronger in competitiveness as 1 − *d*. The standing variation in competitive traits was observed to be almost exactly this value for resources distributed like a power-law (distribution F) for all *d*-values and for *d*-values above the threshold for the other distributions. Thus, standing genetic variation was only comparatively high when strong temporal fluctuations were absent.

**Figure 4.**
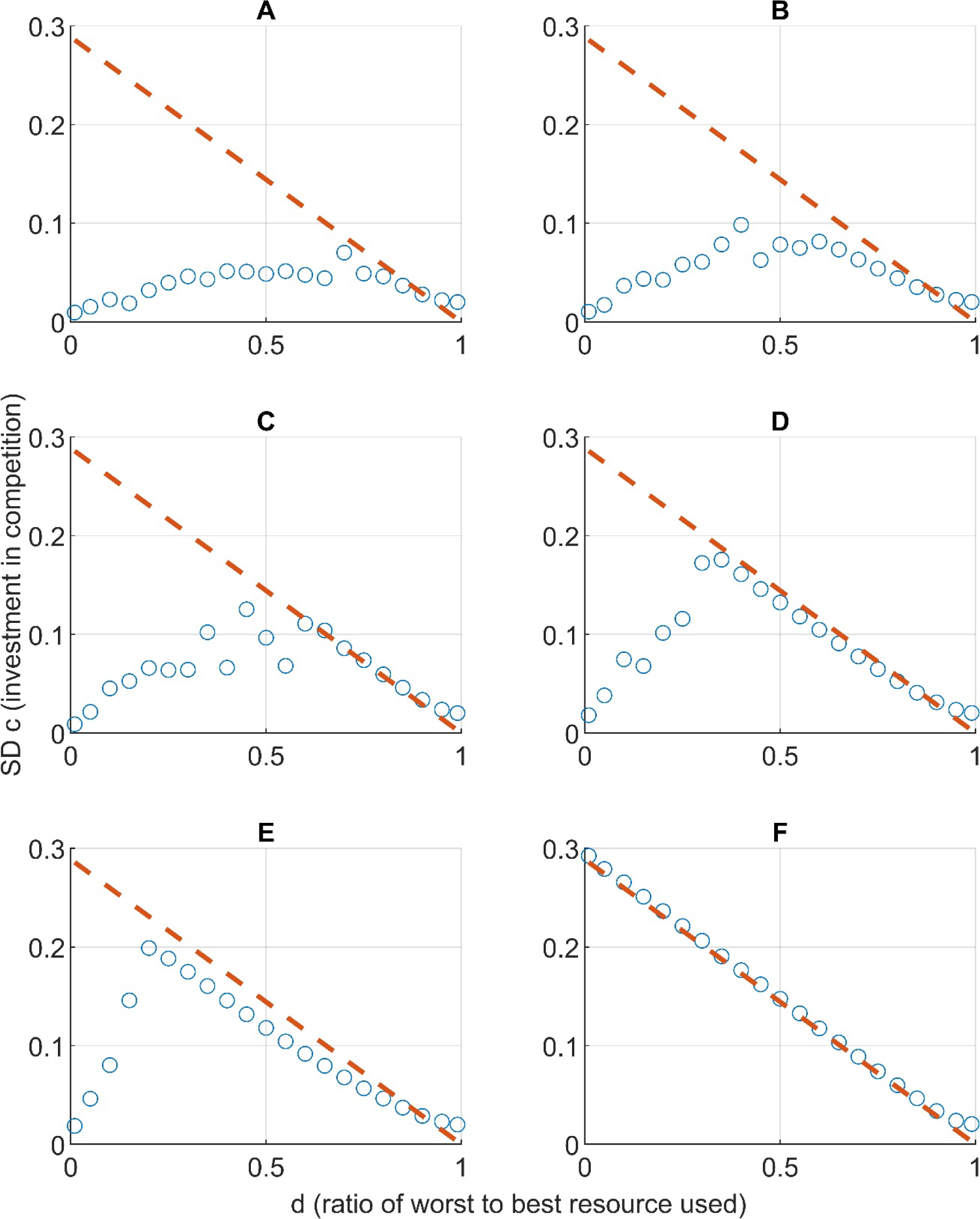
Summary of simulation results dependent on the assumed resource distributions A to F and d, the assumed ratio between the worst and the best resource. Depicted are as an estimate of the extent of maintained genetic variation, the average standard deviations of the competitive trait c, at generation 1000, calculated from 100 independent replicates. The red dashed lines indicate the maximum expected standard deviation SD=(1-d)/(√12).

A closer look reveals some finer details in which the results differ between the various distributions. For a large range of *d* values, the distributions that contain frequent poor and rare good resources, i.e. distributions D to F, show larger standing variation in *c* values at generation 1000, but lower temporal variation in mean *c* values compared to distributions A to C. This is most evident for intermediate values of *d*. This means that distributions with few very good resources resulted in high variation in the competitive trait being selectively maintained in an almost stable fashion for many *d* values. On the other hand, the resource distributions with larger proportions of good resources resulted in strong temporal variation and comparatively low standing variation in the competitive trait at specific timepoints for a broader range of *d* values.

### The special case of the power-law distribution

Distribution F led to results that were divergent from the other distributions. The observed distribution of trait values was, for all *d*-values smaller than 1, indeed approximately uniform, as suggested by the analytical result (see appendix) mentioned earlier. In addition, quick and strong temporal fluctuations in *c*, which were typical for the other resource distributions, did not occur for any *d*-values. The mean *c*-value for this resource distribution was always very close to the predicted value of *d*/2, as expected for a uniform distribution of *c*-values below 1 − *d*.

## Discussion

As expected, our simulations show that larger variation in resource values, i.e. assuming small values of *d*, generally resulted in the evolution of larger investments into the competitive trait *c*. We found that the shape of the resource distribution was also important, in addition to the value of *d*, confirming our hypothesis. Only some resource distributions led to strong temporal variation in the mean value of the competitive trait. At least under some *d* values, all resource distributions led to the maintenance of standing genetic variation in the competitive trait, though the extent of the maintained variation differed strongly between resource distributions. In the following paragraphs, we will discuss how the variation in the mean value of *c*, in the occurrence of temporal fluctuations and in the maintenance of standing variation can be explained.

The ratio between the worst and the best resource available, i.e., the value of *d*, determined to a large extent how much individuals invested on average into the competitive trait. For three vastly different linear distributions (A to C), the average mean trait values we observed were very similar but depended strongly on *d*. In all these cases, the observed average competitive trait *c* was close to the assumed value of 1 − *d*, i.e., the diagonal in the figures, but mean traits tended to be slightly below this value. When good resources were rare, and poor resources were frequent (distributions D and E), mean *c* was similar to that observed under distributions A to C for low values of *d*, but mean *c* decreased more steeply at higher values of *d*. At least for those distributions that include a decent share of high-quality resources, the ratio between the worst and the best resource seems to predict quite well how much individuals evolve to invest on average into competition. This statement is also valid for distribution F, but the relationship is a different one for this distribution, as the average values we observed were closer to (1 − *d*)/2. This result exactly matches our prediction, as this average value is expected for a uniform distribution of trait values between 0 and 1 − *d*. Our study thus seems to give an answer to the question of how much individuals should invest into competition when competition is costly, and when traits can be expected to vary continuously. This is an important novel finding, as, to the best of our knowledge, no other studies have previously addressed this question.

Most resource distributions (A to E) did not lead to a stable mean competitive trait but led to some fluctuations in the mean competitive trait value over time, if *d* was larger than 0 and sufficiently smaller than 1. Thus, under some specific conditions, temporal fluctuations in *c* could be observed. In these cases, adaptive evolution causes cycles of arms races where increasingly higher values of *c* are favoured, but only to a certain point, after which much less competitive mutants can invade. This creates an ‘engine’ that maintains temporal variability in *c*. Strong temporal trait fluctuations resulting from selection as reported by Baldauf et al. (2014) are thus not restricted to the very idealized resource distributions used in that study, but on the other hand strong temporal fluctuations were also absent under many conditions. Strong temporal fluctuations were absent for resource distribution F (independent of the chosen d), as well as for all other examined distributions (A to E) if sufficiently large values of *d* were assumed. In these cases, individuals with low competitive traits on average achieved similar fitness, then those individuals that invested more into competitive traits, leading to an absence of arms races.

Some maintenance of standing genetic variation in competitive trait values was evident in all our simulations, but overall, it was highest when resource distribution F was assumed, and also when poor resources were common (distributions C to E), given that *d* was high enough to prevent strong temporal fluctuations (see figure 4). When the variation maintained was very low, it was likely caused by mutation selection balance. This occurred first when *d* was approaching 1, leading to the evolution of non-competitive trait values, or second, when weak competitors gained very few resources, leading to an arms race for strongly competitive traits. Our results thus show that costly competition can maintain large and continuous genetic and phenotypic variation under very diverse conditions regarding the resource distribution. This is an important new finding. It provides a potential explanation for continuous variation observed, for example, in studies examining individual differences or personality traits (Vukasovic & Bratko 2015). It also potentially explains the surprisingly high values of heritability frequently observed for traits under strong selection (Lynch & Walsh 1998) and is a sufficient condition to explain individualized niches (Krüger et al. 2021, Trappes et al. 2022). As we have used no sexual reproduction or recombination in our model, our results could also be interpreted to show that many species competing for the same resource can coexist. We show that a single resource of variable quality could, for some resource distributions, sustain an arbitrary number of species that make different investments into competition. This conclusion thus contradicts the classical competitive exclusion principle (Hardin 1960).

Our results show that the striking differences in the outcome of two otherwise comparable models (Baldauf et al. 2014, and Kikuchi & Reinhold 2021) likely can be explained by differences in the assumed resource distributions. For some distributions we observed strong temporal fluctuations in the competitive trait for a broad interval of *d*-values, for others, such fluctuations could only be observed for a limited range of values of *d* or could not be observed at all. In line with our results, strong temporal fluctuations are expected for uniform distributions as reported by Baldauf et al. (2014) for *d* = 0.11 and *d* = 0.67, and not expected for distribution E and *d* ≥ 0.25, as used by Kikuchi & Reinhold (2021).

It is also important to consider the limitations of our modelling approach. Most significantly, we employed a model of asexual reproduction. In our model, single individuals reproduced and trait values of offspring were determined by those of their single parents, up to mutations. We did not consider sexual reproduction processes and associated dynamics that can be affected by competitiveness, such as mate choice (Baldauf et al. 2014). However, we note that modelling of evolution that explicitly considers combinations of gametes and other associated dynamics also has a rich legacy in theoretical studies of evolution (DeAngelis & Mooij 2005, Brown & Thomson 2018). Testing if, or to what extent the results in this paper carry over to sexually reproducing organisms is therefore an important aspect of future work. A different important direction of future modelling efforts could be the relaxation of our assumption that the most competitive individual always attains the highest quality resource and so on. This strict hierarchy is a suitable description if individuals can compete with all others independent of location or distance in between. For example, this is the case in well-mixed populations, or systems in which competitive interactions occur over long spatial scales. However, it is not applicable to systems in which competitive (and other) interactions are underpinned by population structure (Perc et al. 2013) such as microbial biofilms (Eigentler et al. 2022) or self-organised vegetation (Gandhi et al. 2019).

It might seem difficult to relate the results of our model to empirical data, for example because the distribution of resources in empirical systems is difficult to measure and to generalize due to the high number of niche dimensions a species may utilize (e.g., Clark et al. 2010). However, the obtained data fit very well with our knowledge about variation of extravagant traits in mate choice and intense intrasexual fighting where losers in competition obtain very few mating opportunities. In line with the evolution of extravagant fighting abilities and highly skewed reproductive success as observed in sea elephants or stag beetles, our model would predict high investment into these competitive traits, as competitors with non-competitive traits in these systems are likely to obtain very low fitness. As we observed the maintenance of large trait variation when exceptionally good resources are comparatively rare so that weak competitors can obtain a share, we predict that larger genetic variance can be maintained. One important prediction of our model would be that different phenotypes can achieve similar fitness values. Future empirical studies should test whether this is the case and whether the genetic variation maintained in competitive traits and the average trait size depend on the resource distributions present. Such resources could for example be represented by territories and their quality or area, groups of mating partners over which the other sex fights, specific spots on a mating lek, or the size of a sexual ornament. When a new habitat is colonized, and population density is low, most individuals likely can achieve such high-quality resources, equivalent to a comparatively high value of *d*. Under such conditions, our model would predict less investment into competition and a high maintenance of genetic variation.

Based on the observed simulation results we conclude that dependent on the present resource distributions, dynamic fluctuations of trait values can but need not occur regularly with costly competition. Such instability has been predicted to be often the outcome of evolutionary games (Nowak & Sigmund 2004) but was absent in our models for resource distribution F, and for distributions A to E if *d* was sufficiently large. Under these conditions, large continuous genetic variation is maintained by negative frequency dependent selection. Stable high investments in competitive traits only developed if weak competitors obtained relatively little fitness. Stable low investment into competitive traits only evolved when very little variation in resource values was present. Our results thus propose explanations for the maintenance of continuous phenotypic variation and the occurrence of temporal fluctuations when individuals compete with costly traits.

## Data availability statement

Modelling code and data have been published in a Github repository and an image at time of submission has been achieved through Zenodo (Reinhold et al. 2023).

## Appendix

## Distribution F leads to uniform trait distributions

In this section of the appendix, we show analytically that for a large number of individuals, the power law resource distribution (F) leads to a stable trait distribution that is uniform, i.e. a distribution in which all trait values 0 ≤ *c* ≤ 1 − *d* occur with the same frequency.

For this, note that the probability density function of resource distribution F is 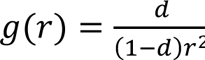. Therefore, its cumulative distribution function is 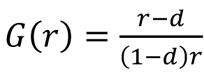. We look for a trait distribution that is stable. In this context, stable corresponds to a setting in which every single individual attains the same fitness, that is the quantity (1 − *c*)*r*(*c*) is constant for all 0 ≤ *c* ≤ 1 − *d*. Since individuals with no investment into competitiveness (*c* = 0) obtain a resource of quality *r*(0) = *d*, we obtain that (1 − *c*)*r*(*c*) = *d* for all individuals with 0 ≤ *c* ≤ 1 − *d*. Rearranging this equality gives 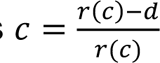.

Due to the hierarchical assignment of resource values *r*(*c*) in response to the trait values *c* present in the population, we can relate the resource distribution with the stable trait distribution through *P*(*r*^∗^ < *r* < 1) = *P*(*c*^∗^ < *c* < 1 − *d*), with *d* < *r*^∗^ < 1, 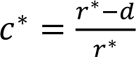.

That is, for any resource value *d* < *r*^∗^ < 1, the number of individuals that obtain higher resource values equals the number of individuals with investment into competition larger than 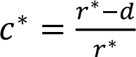 (Fig. A1).

This enables us to obtain the cumulative distribution function of the stable trait distribution through

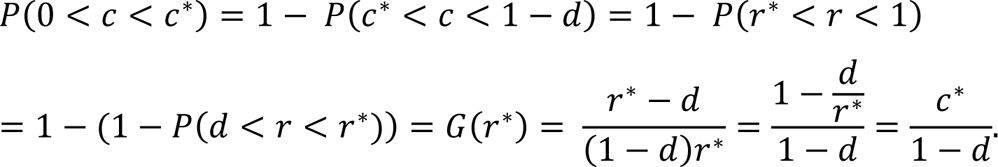

This is the cumulative distribution function of the uniform distribution on 0 ≤ *c* ≤ 1 − *d*. Thus, the power law resource distribution F leads to a uniform trait distribution.

**Figure A1:**
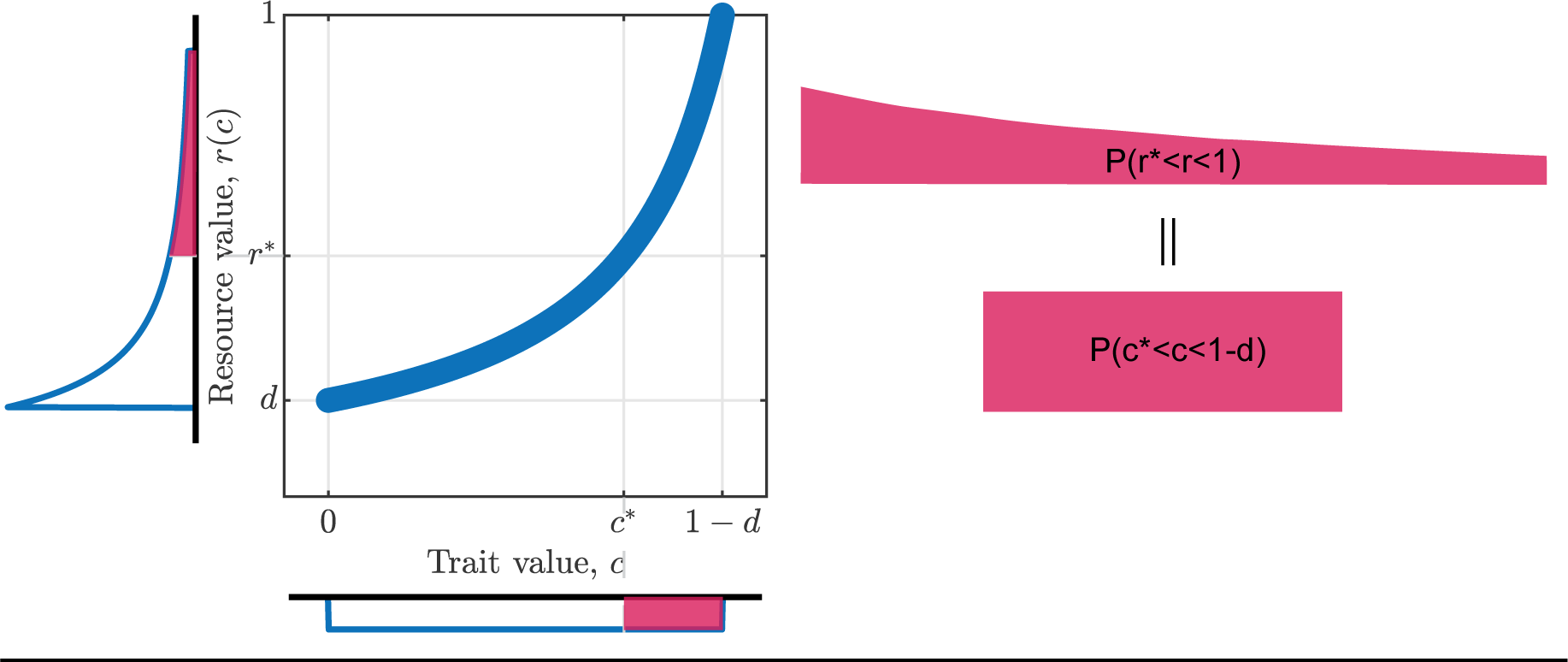
This sketch visualises the relation between trait values and assigned resource values if all individuals attain identical fitness (1 − *c*)*r*(*c*) = *d*. The distribution plot next to the y-axis shows the resource distribution (F). The area of the shaded region is the relative abundance of individuals which attain resource values higher than *r*^∗^. As our analysis shows, this equals the area shaded in the distribution plot for the trait value below the x-axis, which indicates the relative abundance of individuals with investment into competition higher than *c*^∗^ only if the trait distribution is uniform.

## Notes

### Competing Interest Statement

The authors have declared no competing interest.

## References

Baldauf, S.A., Engqvist, L., Weissing, F.J. (2014) Diversifying evolution of competitiveness. Nat. Commun. 5, 5233 (doi: 10.1038/ncomms6233).

Broom, M., Johanis, M., Rychtar, J. (2018) The effect of fight cost structure on fighting behaviour involving simultaneous decisions and variable investment levels. J. Math. Biol. 76, 457–482 (doi: 10.1007/s00285-017-1149-y)

Brown, J.M., Thomson, R.C. (2018) Evaluating model performance in evolutionary biology. Annual Review of Ecology, Evolution, and Systematics 49:95–114 (doi: 10.1146/annurev-ecolsys-110617-062249

Christie, M.R., McNickle, G.G., French, R.A., Blouin, M.S. (2018) Life history variation is maintained by fitness trade-offs and negative frequency-dependent selection. Proc. Natl. Acad. Sci. USA 115, 4441–4446 (doi:10.1073/pnas.1801779115)

Clark, J.S, Bell, D., Chu, C., Courbaud, B., Dietze, M., Hersh, M., HilleRisLambers, J., Ibášez, I., LaDeau, S., McMahon, S., et al. (2010) High-dimensional coexistence based on individual variation: a synthesis of evidence. Ecological Monographs 80, 569–608 (doi:10.2307/20787450)

DeAngelis, D.L., Mooij, W.M. (2005) Individual-based modeling of ecological and evolutionary processes. Annual Review of Ecology, Evolution, and Systematics 36:147–168 (doi: 10.1146/annurev.ecolsys.36.102003.152644)

Eigentler, L., Davidson, F.A., Stanley-Wall N.R. (2022) Mechanisms driving spatial distribution of residents in colony biofilms: an interdisciplinary perspective. Open Biol.12220194220194 (doi: 10.1098/rsob.220194)

Gandhi, P., Iams, S., Bonetti, S., Silber, M. (2019). Vegetation pattern formation in drylands. In: D’Odorico, P., Porporato, A., Wilkinson Runyan, C. (eds) Dryland ecohydrology. Springer, Cham. (doi: 10.1007/978-3-030-23269-6_18)

Grafen, A. (2015) Biological fitness and the fundamental theorem of natural selection. Am. Nat. 186, 1–14 (doi: 10.1086/681585)

Hamilton, W.D., Zuk, M. (1982) Heritable true fitness and bright birds: a role for parasites? Science 218, 384–387

Hardin G. (1960) The competitive exclusion principle. Science 131, 1292–1297. (doi:10.1126/science.131.3409.1292)

Harris, W.E., McKane, A.J., Wolf, J.B. (2008) The maintenance of heritable variation through social competition. Evolution 62, 337–347 (doi:10.1111/j.1558-5646.2007.00302.x)

Houle, D. (1992) Comparing evolvability and variability of quantitative traits. Genetics 130, 195–204 (doi: 10.1093/genetics/130.1.195)

Kikuchi, D.W., Reinhold, K. (2021) Modelling migration in birds: competition’s role in maintaining individual variation. Proc. R. Soc. B. 288, 20210323 (doi: 10.1098/rspb.2021.0323)

Krüger, O., Anaya-Rojas, J., Caspers, B., Chakarov, N, Elliott-Graves, A, Fricke, C. et al. (2021) Individualised niches: an integrative conceptual framework across behaviour, ecology, and evolution. EcoEvoRxiv

Lynch, M., Walsh, B. (1998) Genetics and Analysis of Quantitative Characters. Sunderland, MA: Sinauer

McElreath, R., Strimling, P. (2006) How noisy information and individual asymmetries can make ‘personality’ an adaptation: a simple model. Anim. Behav. 72, 1135–1139

Merilä, J., Sheldon, B.C. (2000) Lifetime reproductive success and heritability in nature. Am. Nat. 155, 301–310 (doi: 10.1086/303330)

Nowak, N.A., Sigmund, K. (2004) Evolutionary dynamics of biological games. Science 303, 793–799 (doi: 10.1126/science.1093411)

Perc, M., Gómez-Gardeñes, J., Szolnoki, A., Floría, L., Moreno, Y. (2013). Evolutionary dynamics of group interactions on structured populations: A review. J R Soc Interface 10: 20120997 (doi:_ 10. 20120997. 10.1098/rsif.2012.0997)

Radwan, J., Engqvist, L., Reinhold, K. (2016) A paradox of genetic variance in epigamic traits: beyond “good genes” view of sexual selection. Evol. Biol. 43, 267–275 (doi: 10.1007/s11692-015-9359-y)

Reinhold, K. (2001) Maintenance of a genetic polymorphism by fluctuating selection on sex-limited traits. J. Evol. Biol. 13, 1009–1014 (doi: 10.1046/j.1420-9101.2000.00229.x)

Reinhold K., Eigentler L., Kikuchi D.W. (2023) Code repository for Reinhold et al.

Evolution of individual variation in a competitive trait: a theoretical analysis (v1.3). Zenodo. (doi: 10.5281/zenodo.8288778)

Sinervo, B., Lively, C.M. (1996) The rock–paper–scissors game and the evolution of alternative male strategies. Nature 380, 240–243

Svardal, H., Rueffler, C., Hermisson, J. (2015) A general condition for adaptive genetic polymorphism in temporally and spatially heterogeneous environments. Theor. Pop. Biol. 99, 76–97 (doi: 10.1016/j.tpb.2014.11.002)

Trappes, R., Nematipour, B., Kaiser, M.I., Krohs, U., van Benthem, K.J., Ernst, et al. (2022) How individualized niches arise: defining mechanisms of niche construction, niche choice, and niche conformance. BioScience 72, 538–548 (doi: 10.1093/biosci/biac023)

Trotter, M.V., Spencer, H.G. (2007) Frequency-dependent selection and the maintenance of genetic variation: exploring the parameter space of the multiallelic pairwise interaction model. Genetics 176, 1729–1740 (doi: 10.1534/genetics.107.073072)

Vukasović, T., Bratko, D. (2015) Heritability of personality: A meta-analysis of behavior genetic studies. Psychol. Bull.141:769–785 (doi: 10.1037/bul0000017)

Wolf, M., van Doorn, G.S., Leimar, O., Weissing, F.J. (2007) Life-history trade-offs favour the evolution of animal personalities. Nature 447, 581–584 (doi:10.1038/nature05835)

Wolf, M., van Doorn, G.S., Weissing, F.J. (2008) Evolutionary emergence of responsive and unresponsive personalities. Proc. Natl Acad. Sci. USA 105, 15 825–15 830 (doi:10.1073/pnas.0805473105)

